# BNIP3 phosphorylation by JNK1/2 promotes mitophagy via enhancing its stability under hypoxia

**DOI:** 10.1101/2020.08.27.271270

**Authors:** Yun-Ling He, Sheng-Hui Gong, Xiang Cheng, Ming Zhao, Tong Zhao, Yong-Qi Zhao, Ming Fan, Ling-Ling Zhu, Li-Ying Wu

**Affiliations:** Department of Cognitive Sciences, Institute of Cognition and Brain Sciences, Beijing, 100850, China; Co-Innovation Center of Neuroregeneration, Nantong University, Nantong, 226001, China; Beijing Institute for Brain Disorder, Beijing, 102206, China; State Key Laboratory of Proteomics, Beijing Proteome Research Center, Beijing Institute of Radiation Medicine, Beijing, 100850, China

**Keywords:** BNIP3, hypoxia, mitophagy, phosphorylation, ubiquitination

## Abstract

Mitophagy is an important metabolic mechanism that modulates mitochondrial quality and quantity by selectively removing damaged or unwanted mitochondria. BNIP3, a mitochondrial outer membrane protein, is a mitophagy receptor that mediates mitophagy under various stresses, particularly hypoxia, since BNIP3 is a hypoxia-responsive protein. However, the underlying mechanisms that regulate BNIP3 and thus mediate mitophagy under hypoxic conditions remain elusive. Here, we demonstrate that in hypoxia JNK1/2 phosphorylates BNIP3 at Ser 60/Thr 66, which hampers proteasomal degradation of BNIP3 and drives mitophagy by facilitating the direct binding of BNIP3 to LC3, while PP1/2A represses mitophagy by dephosphorylating BNIP3 and triggering its proteasomal degradation. These findings reveal the intrinsic mechanisms cells use to regulate mitophagy via the JNK1/2-BNIP3 pathway in response to hypoxia. Thus, the JNK1/2-BNIP3 signaling pathway strongly links mitophagy to hypoxia and may be a promising therapeutic target for hypoxia-related diseases.

## Introduction

Under normal circumstances, functional mitochondria are energy factories that provide the cellular ATP required for cellular activities. However, under hypoxic conditions, mitochondria become sites where excessive reactive oxygen species (ROS) are generated, which in turn impairs mitochondria function (Scherz-Shouval & Elazar, 2011). The damaged mitochondria release proteins that participate in the initiation of apoptosis (Ashrafi & Schwarz, 2013; Kubli & Gustafsson, 2012; Marino *et al*, 2014). To defend against the harmful effects of dysfunctional mitochondria and maintain homeostasis, cells initiate protective mechanisms to compensate for damaged mitochondria prior to suffering harm. Mitophagy is recognized as a major protective mechanism by which dysfunctional mitochondria are cleared to enhance overall mitochondria quality and simultaneously provide rapid recycling of metabolites (Palikaras *et al*, 2018). The mitophagy process is complicated and involves mitochondrial dynamics, recognition and labeling of target mitochondria, envelopment of mitochondria by autophagosomes, fusion of autophagosomes-lysosomes and degradation of mitochondria by proteases in lysosomes (Ashrafi & Schwarz, 2013; Palikaras *et al*., 2018; Youle & Narendra, 2011). Among these processes, recognizing and labeling damaged or unwanted mitochondria is a critical process, considering that mitophagy is one of selective autophagy.

Two critical recognition signals are required for mitophagy in mammalian cells: ubiquitin (Ub)-adaptor- and receptor-mediated mitochondrial conjunction with LC3 (microtubule-associated protein 1 light chain 3) on the autophagosomes (Dikic & Elazar, 2018). Ubiquitin-adaptor-mediated mitophagy is provoked by the classic PINK1 (PTEN-induced kinase 1) and E3 ubiquitin protein ligase Parkin pathway (Pickrell & Youle, 2015), and receptor-mediated mitophagy is initiated by several mitochondrial outer membrane proteins: the BH3-only proteins BNIP3 (BCL2/adenovirus e1B 19 kDa protein interacting protein 3) and BNIP3L (BNIP3-like, also known as NIX), which are 56% identical (Hamacher-Brady & Brady, 2016; Kubli & Gustafsson, 2012), FUNDC1 (FUN14 domain-containing protein 1), etc. (Kirkin & Rogov, 2019; Liu *et al*, 2014) In the former signaling pathway, when the mitochondrial membrane potential is lost under stress conditions, PINK1, a mitochondrial kinase, accumulates on the outer membrane of mitochondria (Narendra *et al*, 2010) and activates the E3 ubiquitin ligase activity of Parkin via phosphorylation of Parkin and ubiquitin (Kane *et al*, 2014; Koyano *et al*, 2014), which recruits Parkin to the damaged mitochondria (Geisler *et al*, 2010; Narendra *et al*., 2010). Parkin selectively ubiquitinates its substrates on the outer membrane of mitochondria, and the ubiquitin-conjugated substrates are recognized by ubiquitin-binding adaptor proteins (for example p62, NDP52, and OPTN) that link with LC3 on the developing autophagosomes, leading to mitochondrial sequestration by the autophagosomes and removal by the lysosomes (Geisler *et al*., 2010; Heo *et al*, 2015; Lazarou *et al*, 2015). In the latter signaling pathway, previous studies have shown that mitophagy receptors possess the characteristic recognition sequence W/F/YxxL/I/V for LC3 that mediates selective autophagy (Rogov *et al*, 2014). Coincidentally, many mitophagy receptors are responsive to hypoxia. A series of reports on FUNDC1 indicate that under normoxic conditions, FUNDC1 is phosphorylated at Tyr 18 and Ser 13 by Src and CK2 (casein kinase 2), which hampers the interaction of LC3 with FUNDC1; under hypoxic conditions or loss of mitochondrial membrane potential, Src and CK2 are unable and ULK1 (Unc-51-like kinase 1) is able to phosphorylate FUNDC1 at Ser 17, and PGAM5 (phosphoglycerate mutase family member 5) dephosphorylates FUNDC1 at Ser 13, all of which lead to enhanced interaction of FUNDC1 and LC3, and hence, mitophagy induction (Chen *et al*, 2014; Liu *et al*, 2012; Wu *et al*, 2014). To date, the regulation mechanisms of FUNDC1- and PINK1/Parkin-mediated mitophagy have been well illustrated. However, until now, the mechanisms underlying BNIP3/BNIP3L-mediated mitophagy, especially under hypoxic conditions, have been far from clear. Although it has been demonstrated that the phosphorylation of BNIP3 at Ser 17/24 and BNIP3L at Ser 34/35 or Ser 81 enhances their respective association with LC3 and facilitates activation of mitophagy (Rogov *et al*, 2017; Yuan *et al*, 2017; Zhu *et al*, 2013), but the kinases and phosphatases targeting BNIP3/BNIP3L have not yet been uncovered.

BNIP3 is a member of the atypical BH3-only subfamily within the BCL-2 family and is localized at the mitochondrial outer membrane (Chen *et al*, 1997; Yasuda *et al*, 1998). BNIP3 is transcriptionally activated by the transcription factor HIF-1 (hypoxia-inducible factor 1) under hypoxia, which is why it is extremely sensitive to hypoxia and is generally used as a typical target gene of HIF-1 (Bruick, 2000; Guo *et al*, 2001; Sowter *et al*, 2001). In addition, as far as we know, BNIP3 is more sensitive to hypoxia than any of the other proteins on the mitochondrial outer membrane (Bruick, 2000). The functions of BNIP3 appear to be contradictory, involving the induction of apoptosis or mitophagy in different contexts or cell types (Bellot *et al*, 2009; Chourasia *et al*, 2015; Diwan *et al*, 2007; Ney, 2015; Zhang *et al*, 2008); however, the precise mechanisms of BNIP3 functions have not been elucidated. In recent years, many studies have focused on BNIP3-mediated mitophagy, but how BNIP3 regulates mitophagy under hypoxia remains unclear. It has been demonstrated that overexpression of BNIP3 under hypoxic conditions promotes its interaction with BCL-2/BCL-XL, which contributes to the release of Beclin-1 from BCL-2/BCL-XL binding and initiates autophagy (Bellot *et al.*, 2009; Zhang *et al.*, 2008). However, after induction of autophagy by overexpression of BNIP3, how damaged mitochondria are recognized to initiate mitophagy in the same context has not been clarified. Some studies have reported that BNIP3 overexpression promotes mitophagy by enhancing its interaction with the autophagosome membrane protein LC3 (Hanna *et al*, 2012; Ma *et al*, 2012). Here, we demonstrate that BNIP3 phosphorylation, rather than its overexpression, plays a decisive role in mediating mitophagy.

In this study, we identified the new phosphorylation site Ser 60/Thr 66 in BNIP3. Most importantly, we identified JNK1/2 (c-Jun N-terminal kinase 1/2) and PP1/2A (protein phosphatase 1/2A) as the kinase and phosphatase responsible for phosphorylation and dephosphorylation, respectively, of BNIP3 at Ser 60/Thr 66 residue in response to hypoxia. Furthermore, we demonstrated that phosphorylation of BNIP3 by JNK1/2 is required for both induction of mitophagy and increased stability of BNIP3, while dephosphorylation of BNIP3 by PP1/2A causes proteasomal degradation of BNIP3 and accordingly failure of mitophagy induction. To the best of our knowledge, the mechanisms by which BNIP3 is degraded via the ubiquitin-proteasome pathway have not previously been revealed. Here, we report the crosstalk between BNIP3-mediated mitophagy and its proteasomal degradation under hypoxic conditions. Collectively, our study shows that JNK1/2 and PP1/2A oppositely regulate BNIP3 phosphorylation and consequently manipulate its stability, which in turn affects the induction of mitophagy.

## Results

### Phosphorylation of BNIP3 is related to mitophagy under hypoxia

We first observed mitophagy under different hypoxia conditions using an oxygen-sensitive PC12 cell line (Millhorn *et al*, 1996), and surprisingly found that mitophagy under different hypoxia conditions is vastly different. Compared with normoxia (20% O_2_), 10% O_2_ promoted mitophagy and 0.3% O_2_ suppressed mitophagy when cells were exposed to the different oxygen levels for the same time, based on analysis of the morphological characteristics of mitochondria via transmission electron microscopy (TEM) and TOMM20 expression, and the co-localization of autophagosomes and mitochondria via a fluorescence confocal microscopy (Fig. 1A, B and Fig. EV1A). To investigate the possible causes of different mitophagy under different hypoxia conditions, we then evaluated the effects of BNIP3 on mitophagy under different hypoxia, because BNIP3 is extremely sensitive to hypoxia stimulation except as a mitophagy receptor. Notably, we found that the BNIP3 protein bands in the 10% O_2_ group lay at the top (30 kDa), while most of the bands in the 0.3% O_2_ group accumulated beneath the top 24 h after exposure to hypoxia. Comparatively, the protein level at the top was higher in the 10% O_2_ group than in the 0.3% O_2_ group but not the total protein level. Meanwhile, we noticed that the BNIP3 protein level at the top was consistent with the mitophagy activity (Fig. 1C). In addition, when *Bnip3* was knocked down with siRNA, mitophagy in the 10% O_2_ group was obviously attenuated, with reduced levels of the autophagosome marker LC3-II and increased levels of the mitochondrial outer membrane protein TOMM20, while knockdown of *Bnip3* in the 0.3% O_2_ group did not lead to comparable differences in mitophagy (Fig. 1D). It such seems that the high BNIP3 protein level in the 0.3% O_2_ group was not conducive to mitophagy activity. Taken together, these data indicate that regulation of mitophagy by BNIP3 is not completely dependent on its protein level.

**Figure 1.**
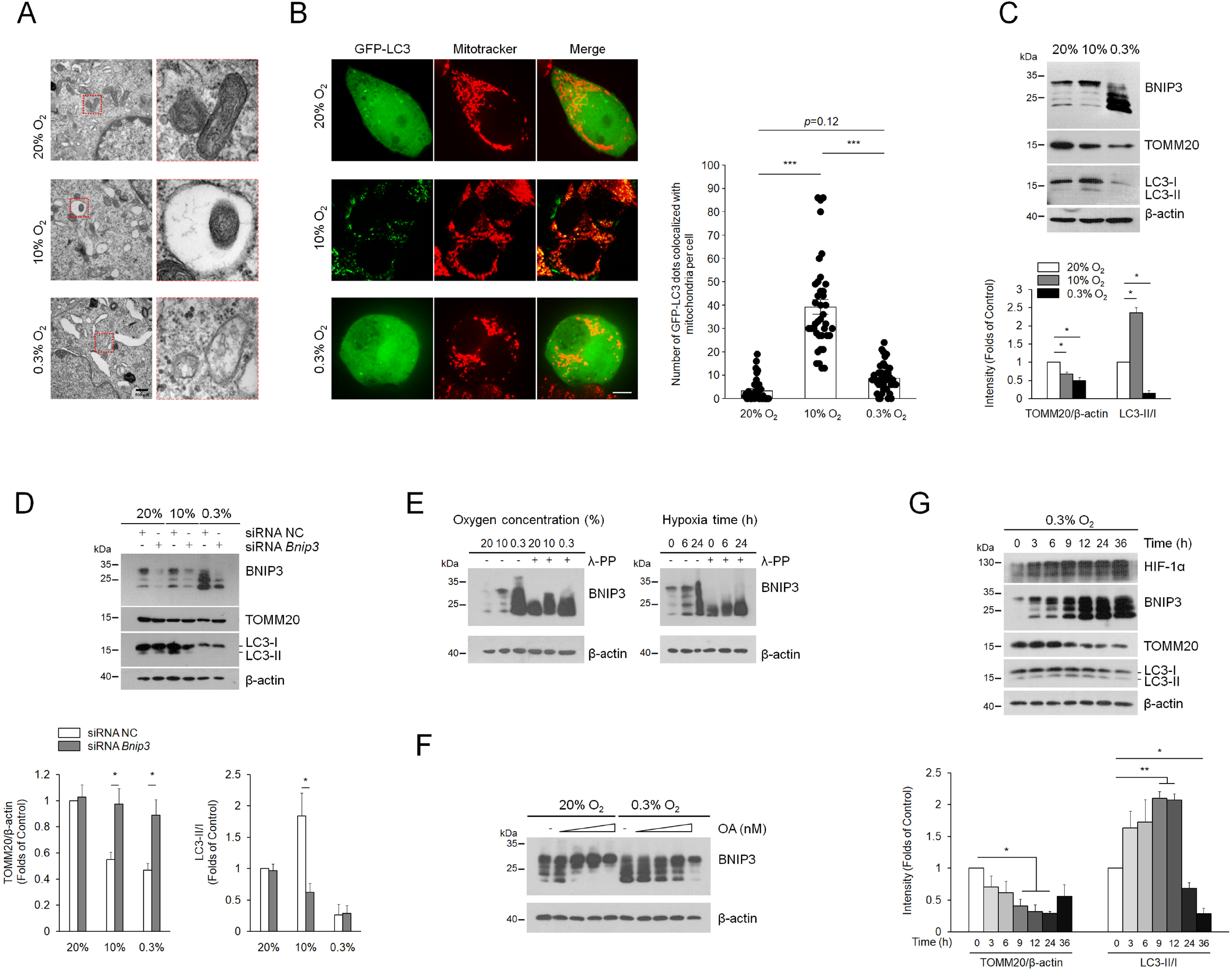
BNIP3 phosphorylation is closely related to mitophagy. A Mitochondria morphology was analyzed via transmission electron microscopy after PC12 cells were exposed to 20% O_2_, 10% O_2_ or 0.3% O_2_ for 24 h. The red boxes indicate representative mitochondria exposed to different oxygen conditions. Scale bar, 500 nm. B PC12 cells were treated the same as in (A), and mitophagy was identified and quantified by co-localization of autophagosomes (GFP-LC3, green) and mitochondria (Mitotracker, red). Scale bar, 10 μm. *n* = 40. C PC12 cells were treated same as (A), BNIP3, TOMM20, LC3 and adaptor protein p62 were detected via western blot. *n* = 3. D PC12 cells were transfected with negative control (NC) or *Bnip3* siRNA for 48 h and then exposed to different oxygen concentrations for 24 h. The levels of BNIP3 mitophagy related proteins were detected via western blot. *n* = 3. E PC12 cells were exposed to different oxygen concentrations for 24 h or 0.3% O_2_ for the indicated times. Cell lysates were treated with/without lambda phosphatase (λ-PPase) for 1 h, followed by western blot analysis. F PC12 cells were exposed to 20% O_2_ or 0.3% O_2_ complemented with 0, 50, 100, 200, 500 nM okadaic acid (OA), and subjected to western blot. G PC12 cells were exposed to 0.3% O_2_ for the indicated times. Cell lysates were then immunoblotted for the indicated proteins. *n* = 3. The data are expressed as means ± SEM. \**P* < 0.05, \*\**P* < 0.01, \*\*\**P* < 0.001 versus the indicated group.

Considering post-translational modifications, we speculated that phosphorylation and dephosphorylation might be involved in regulating the differences in BNIP3 protein bands. To determine if the top BNIP3 band was a result of phosphorylation modification, we added lambda phosphatase (λ-PPase) to cell lysates and excitedly found that all the upper bands migrated downwards at 24 h after exposure to different oxygen concentrations or exposure for different time intervals (Fig. 1E). Conversely, after cells were exposed to the phosphatase inhibitor okadaic acid (OA), nearly all the BNIP3 protein bands in the normoxia (20% O_2_) group quickly accumulated at the top, and the bands in the hypoxia (0.3% O_2_) group gradually migrated upwards and eventually reached the top as the concentration of OA was increased (Fig. 1F). From the above results, we inferred that BNIP3 is regulated by multisite phosphorylation, and the fully phosphorylated form (hereafter referred to as phosphorylation of BNIP3) is located at the top of protein bands, consistent with the description in previous reports that BNIP3 exists in multiple phosphorylated forms (Graham *et al*, 2007; Mellor *et al*, 2010). To clarify whether the BNIP3 phosphorylation contributes to mitophagy induction, we examined the correlation between BNIP3 phosphorylation and mitophagy by comparing the dynamic changes in related proteins under 0.3% O_2_ hypoxic conditions in PC12 cells. The results clearly showed that when the levels of BNIP3 phosphorylation increased with time during early hypoxia, mitophagy was augmented, with increased LC3-II and lower TOMM20 levels; by contrast, when BNIP3 phosphorylation decreased during late hypoxia, mitophagy was suppressed, with less LC3-II and undiminished TOMM20 expression, which was verified in the Hela cells that is used widely for genetic engineering (Fig. 1G and Fig. EV1B). Collectively, these preliminary results demonstrate that BNIP3 phosphorylation is closely associated with mitophagy activation.

### Phosphorylation of BNIP3 at S60/T66 is necessary to promote mitophagy via enhancing its interaction with LC3

To find potential phosphorylation sites in BNIP3, we first searched the “PhosphoSite” database (http://www.phosphosite.org). Based on reported proteomics data, a total of 12 BNIP3 phosphorylation events based on mass spectra evidence are shown in the schematic in Fig. 2A, which represent highly conserved serine or threonine (S/T) residues across vertebrates. To further determine the specific phosphorylation sites that affect the migration of BNIP3 protein bands, mutants were constructed in which each of the 12 S/T residues was changed to alanine (A) via site-directed mutagenesis to inactivate phosphorylation. HeLa cells were transfected with Flag-tagged BNIP3-S/T to A mutants, and the protein phosphorylation was assessed on the basis of changes in the migration of the protein bands after treatment with OA. The results revealed that S/T-to-A replacement at Ser 60 (S60A) blocked the upshift of BNIP3 protein bands (Fig. 2B) and double replacement at Ser 60 and Thr 66 (S60/T66A) potentiated this effect (Fig. 2C), given that these two sites have the common motif recognized by MAPKs (mitogen-activated protein kinases) or CDKs (cyclin-dependent kinases). These data indicate that Ser 60 and Thr 66 are the potential phosphorylation sites of BNIP3 and the former is the primary one, the latter is synergistic. To validate the two phosphorylation sites of BNIP3, we first immunoprecipitated Flag-BNIP3 and tested the precipitates with an antibody against phospho-MAPK/CDK substrates. As expected, both the S60A and S60/T66A mutants reduced the levels of potential phosphorylation of BNIP3 compared to wild-type (WT) (Fig. EV2A). Then, we produced a specific antibody directed to phospho-Ser 60 (p-S60) of BNIP3. This antibody recognized WT, but not the S60A and S60/T66A mutants (Fig. 2D). Using the p-S60 antibody we observed that BNIP3 phosphorylation was increased in the early stage of hypoxia and reduced in the late stage of hypoxia (Fig. 2E). Altogether, these data suggest that Ser 60 is the primary phosphorylation site of BNIP3. Thereafter, we also examined the effects of hypoxia on phosphorylation of BNIP3 at Ser 60 and the effects of disabling phosphorylation at Ser 60/Thr 66 on mitophagy in one experiment. It is clear that hypoxia increased the phosphorylation of BNIP3 at Ser 60 and promoted mitophagy with reduced TOMM20 and increased LC3-II. Introduction of siRNA-resistant phosphorylation-disabled S60/T66A mutant after *Bnip3* knockdown with siRNA impaired the effect of hypoxia on mitophagy (Fig. 2F). These results indicate that the phosphorylation of BNIP3 at Ser 60/Thr 66 is required for promoting mitophagy.

**Figure 2.**
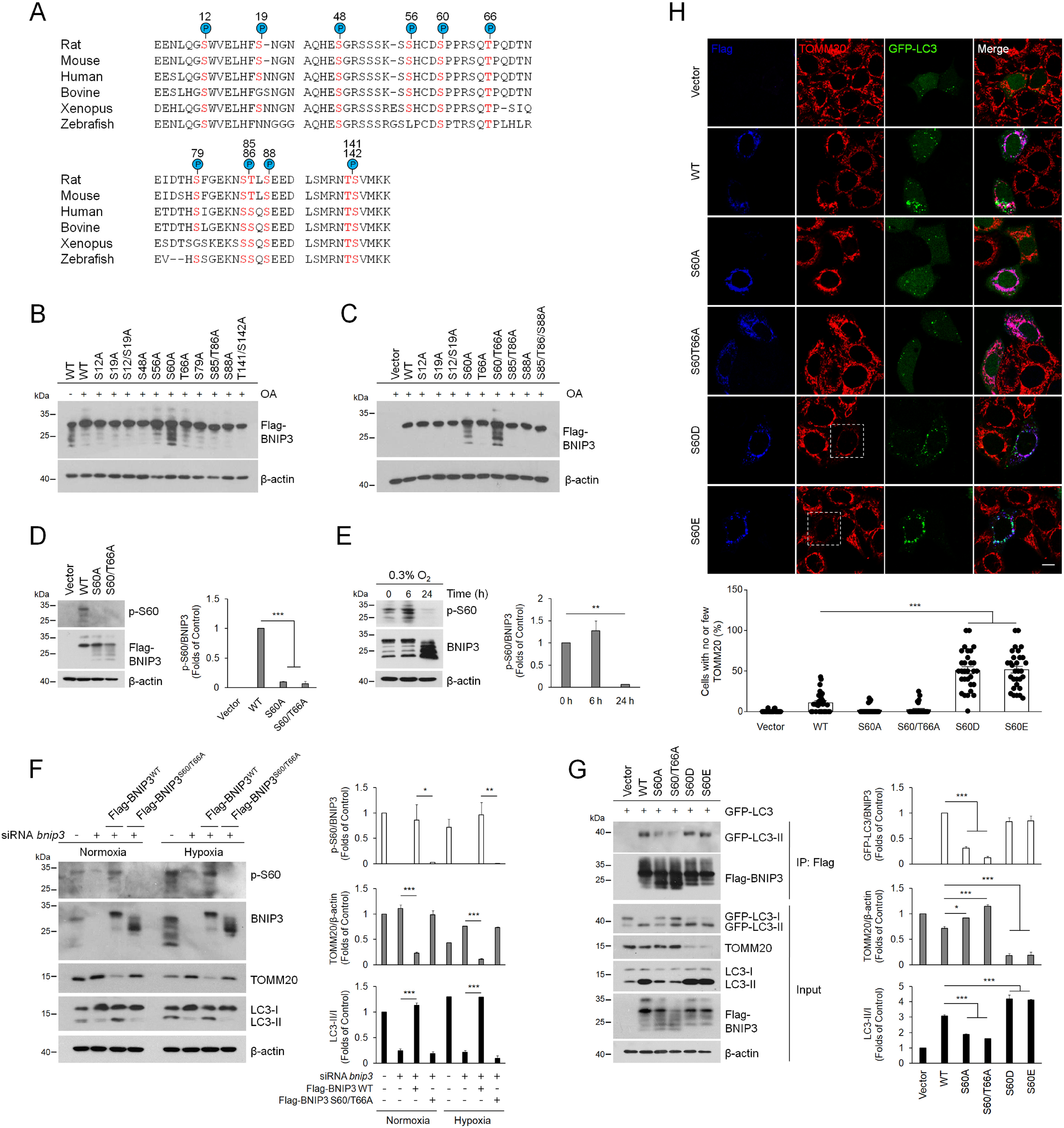
Phosphorylation of BNIP3 at S60/T66 is necessary to promote mitophagy by enhancing its interaction with LC3. A Putative BNIP3 phosphorylation sites based on reported proteomics data (PhosphoSitePlus^®^) that are conserved across different species. Conserved serine and threonine residues are marked in red. B, C HeLa cells were transfected with empty vector and Flag-BNIP3 plasmids encoding either wild-type (WT) or mutant BNIP3 constructs generated via site-directed mutagenesis. After 48 h of transfection, cell lysates were detected via western blotting with anti-Flag antibody. OA, okadaic acid. D HeLa cells were transfected with empty vector, WT, S60A or S60/T66A mutant Flag-BNIP3 plasmids, phosphorylation of BNIP3 was detected via western blotting using a phosphospecific antibody against BNIP3 at Ser 60 (p-S60). *n* = 3. E The phosphorylation level of BNIP3 at Ser 60 in PC12 cells was measured under 0.3% O_2_ for the indicated time. *n* = 3. F PC12 cells were treated with *Bnip3* siRNA and transfected with *Bnip3* siRNA-resistant WT or S60/T66A plasmids, and then cells were exposed to 20% O_2_ (Normoxia) or 0.3% O_2_ (Hypoxia) for 6 h, respectively. Mitophagy and the phosphorylation of BNIP3 were detected via western blotting. *n* = 3. G HeLa cells were transfected with GFP-LC3 and empty vector, WT or the indicated Flag-BNIP3 mutants for 48 h. Cell lysates were immunoprecipitated with anti-Flag antibody and then subjected to western blot analysis with the indicated protein antibodies. *n* = 3. H HeLa cells were treated the same as in (G). The cells were then fixed and immunostained with Flag (blue) and TOMM20 (Red). The white boxes indicate cells with few mitochondria. Scale bars, 10 μm. The data are expressed as means ± SEM. \**P* < 0.05, \*\**P* < 0.01, \*\*\**P* < 0.001 versus the indicated group.

According to previous studies that overexpression of BNIP3 in hypoxia competes with Beclin-1 to interact with BCL-2, followed by release of Beclin-1 and induction of mitophagy (Bellot *et al*., 2009; Zhang *et al*., 2008), we next investigated whether the phosphorylation of BNIP3 at Ser 60/Thr 66 site could affect its association with BCL-2 and thus affect the induction of mitophagy. Regrettably, all mutations of BNIP3 displayed no significant differences in interaction with BCL-2 compared to the WT BNIP3 (Fig. EV2B), suggesting that the phosphorylation of BNIP3 at least at Ser 60/Thr 66 site is not related with the interaction between BNIP3 and BCL-2. After that, we examined whether the phosphorylation of BNIP3 Ser 60/Thr 66 would affect its binding to LC3. Cells expressing GFP-tagged LC3 and Flag-tagged WT or mutant BNIP3 were collected for co-IP assays. In line with previous studies (Hanna *et al*., 2012; Ma *et al*., 2012), GFP-LC3-II was co-precipitated with BNIP3 WT, whereas the interaction was abated by S60A or S60/T66A mutant and mitophagy was inhibited. By contrast, a BNIP3-S60 to aspartic acid (S60D) or to glutamic acid (S60E) mutation that mimics phosphorylation enhanced the binding affinity between BNIP3 and GFP-LC3-II and increased mitophagy (Fig. 2G). To demonstrate further whether the phosphorylation of BNIP3 at Ser 60/Thr 66 impacts mitophagy, we expressed these BNIP3 mutants in cells and detected mitophagy via fluorescence confocal microscopy. We observed that the S60A and S60/T66A mutants led to inhibition of mitophagy, while the S60D and S60E mutants induced more pronounced mitophagy, as shown by the appearance of more LC3 puncta and fewer mitochondria (Fig. 2H). Taken together, these results indicate that phosphorylation of BNIP3 at Ser 60/Thr 66 is necessary for its interaction with LC3 and induction of mitophagy.

### Phosphorylation of BNIP3 at S60/T66 is essential to improve its stability

In addition to the correlation with mitophagy, BNIP3 phosphorylation is also related to its stability. We unexpectedly found that the phosphatase inhibitor OA hindered rapid degradation of BNIP3 when protein synthesis was inhibited with cycloheximide (CHX) under 0.3% O_2_ conditions. The proteasome inhibitor MG132 led to more protein accumulated, but as the concentration of OA increased, the accumulation of BNIP3 by MG132 was reduced (Fig. 3A, B), indicating that BNIP3 phosphorylation may impede its proteasomal degradation. To determine whether phosphorylation at the Ser 60/Thr 66 site is involved in regulation of BNIP3 stability, we measured the effect of the phospho-disabling or phospho-mimic mutations of these sites on BNIP3 degradation after HeLa cells were transfected with the mutants and treated with CHX. Our data demonstrated that the S60A and S60/T66A mutants accelerated the degradation of BNIP3, and the S60D and S60E mutants dramatically hampered this process, suggesting that phosphorylation at Ser 60/Thr 66 is required for BNIP3 stability (Fig. 3C-F). Since ubiquitination usually leads to proteasomal degradation, we further detected the relationship between phosphorylation and ubiquitination of BNIP3. As shown in Fig. 3G, the phospho-disabling and phospho-mimic BNIP3 mutants were linked with more or fewer ubiquitin molecules than the wild-type protein, respectively, which confirmed our above findings. Collectively, these results indicate that phosphorylation of BNIP3 at Ser 60/Thr 66 promotes its stability. Thus, we propose that the improved BNIP3 stability may be the premise and foundation for the induction of mitophagy.

**Figure. 3.**
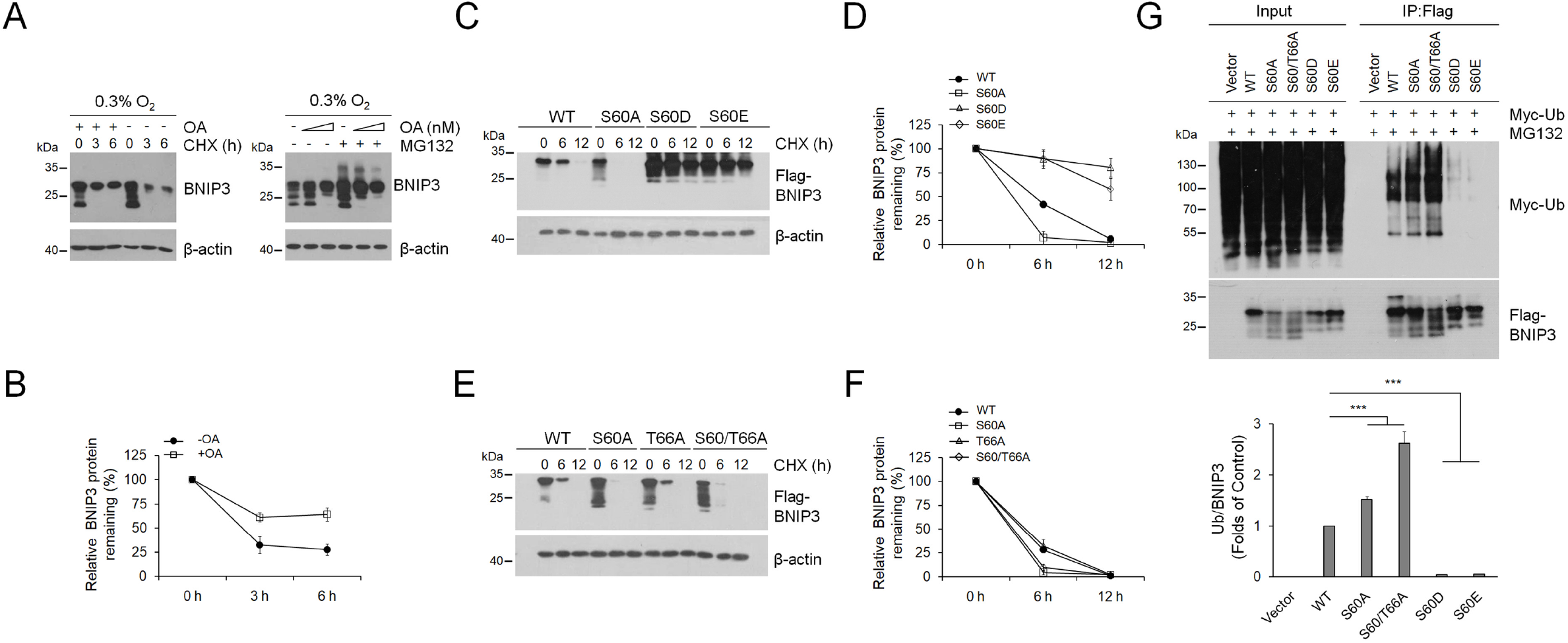
Phosphorylation of BNIP3 at S60/T66 is essential to improve its stability. A PC12 cells were exposed to 0.3% O_2_ supplemented with 200 nM okadaic acid (OA) for 9 h and then treated with 20 μg ml^-1^ cycloheximide (CHX) for the indicated times (left panel) or with 100, 200 nM OA plus 10 μM MG132 for 6 h (right panel). Cell lysates were detected via western blotting using the indicated antibodies. *n* = 3. B Quantification of degradation rate of BNIP3 in CHX chase experiments shown in (A). C-F HeLa cells were transfected with plasmids encoding WT or the indicated BNIP3 mutants for 48 h and then treated with 20 μg ml^-1^ CHX for 0 h, 6 h, or 12 h. BNIP3 expression was detected via western blotting. (D) and (F) are quantification of degradation rate of BNIP3 shown in (C) and (E), respectively. *n* = 3. G HeLa cells were transfected with Myc-Ub and empty vector, WT or the indicated BNIP3 mutant for 48 h and treated with 10 μM MG132 for 12 h. Cell lysates were boiled and immunoprecipitated with an anti-Flag antibody. The immune complexes were then analyzed via western blotting. *n* = 3. The data are expressed as means ± SEM. \**P* < 0.05, \*\**P* < 0.01, \*\*\**P* < 0.001 versus the indicated group.

### JNK1/2 is the kinase responsible for phosphorylation of BNIP3 at S60/T66

Subsequently, we sought to identify which kinases are responsible for the phosphorylation of BNIP3 Ser 60/Thr 66 to better understand the mechanism by which BNIP3 is phosphorylated and thereby mediates mitophagy activation. Since the Ser 60/Thr 66 residue within the consensus motif of MAPKs and CDKs, we screened BNIP3-specific kinases using the respective inhibitors of MAPKs and CDKs. The results showed that the JNK inhibitor SP600125 and MEK1/2 (mitogen-activated protein kinase kinase 1/2) inhibitor PD184352 caused a marked decrease in BNIP3 phosphorylation and an increase in the dephosphorylated forms of BNIP3. Cell cycle-related inhibitors, such as roscovitine (a selective CDK inhibitor), did not significantly affect the features of BNIP3 protein bands, suggesting that BNIP3 is not a phosphorylation substrate of CDKs. Another MAPK inhibitor, SB203580 (a specific p38-MAPK inhibitor), did not have an apparent impact on the BNIP3 protein bands. Additionally, neither K252c nor Bis I (selective PKC inhibitors) nor TBB (a selective CK2 inhibitor) affected the features of BNIP3 protein bands (Fig. 4A), although PKC and CK2 have ever been reported to be related to phosphorylation of BNIP3 or other mitophagy receptors (Chen *et al*., 2014; Graham *et al*., 2007; Kanki *et al*, 2013; Zhu *et al*., 2013). We then combined SP600125 or PD184352 and OA to treat cells and observed that the upshift of BNIP3 protein bands caused by OA was partly reversed by SP600125 but not affected by PD184352 (Fig. EV3A). Moreover, two different JNK inhibitors, SP600125 and JNK-IN-8, could reduce BNIP3 phosphorylation under normoxia or hypoxia (Fig. 4B). The above results suggest that JNK may be the potential kinase for BNIP3. We further determined the effect of SP600125 on the phosphorylation of BNIP3 at Ser 60/Thr 66 in cells expressing WT or S60/T66A mutant. Consistent with changes of BNIP3 bands, the phosphorylation of WT but not S60/T66A mutant BNIP3 was affected by SP600125 (Fig. 4C), suggesting that the Ser 60/Thr 66 residue in BNIP3 may be the target site for JNK recognition.

**Figure 4.**
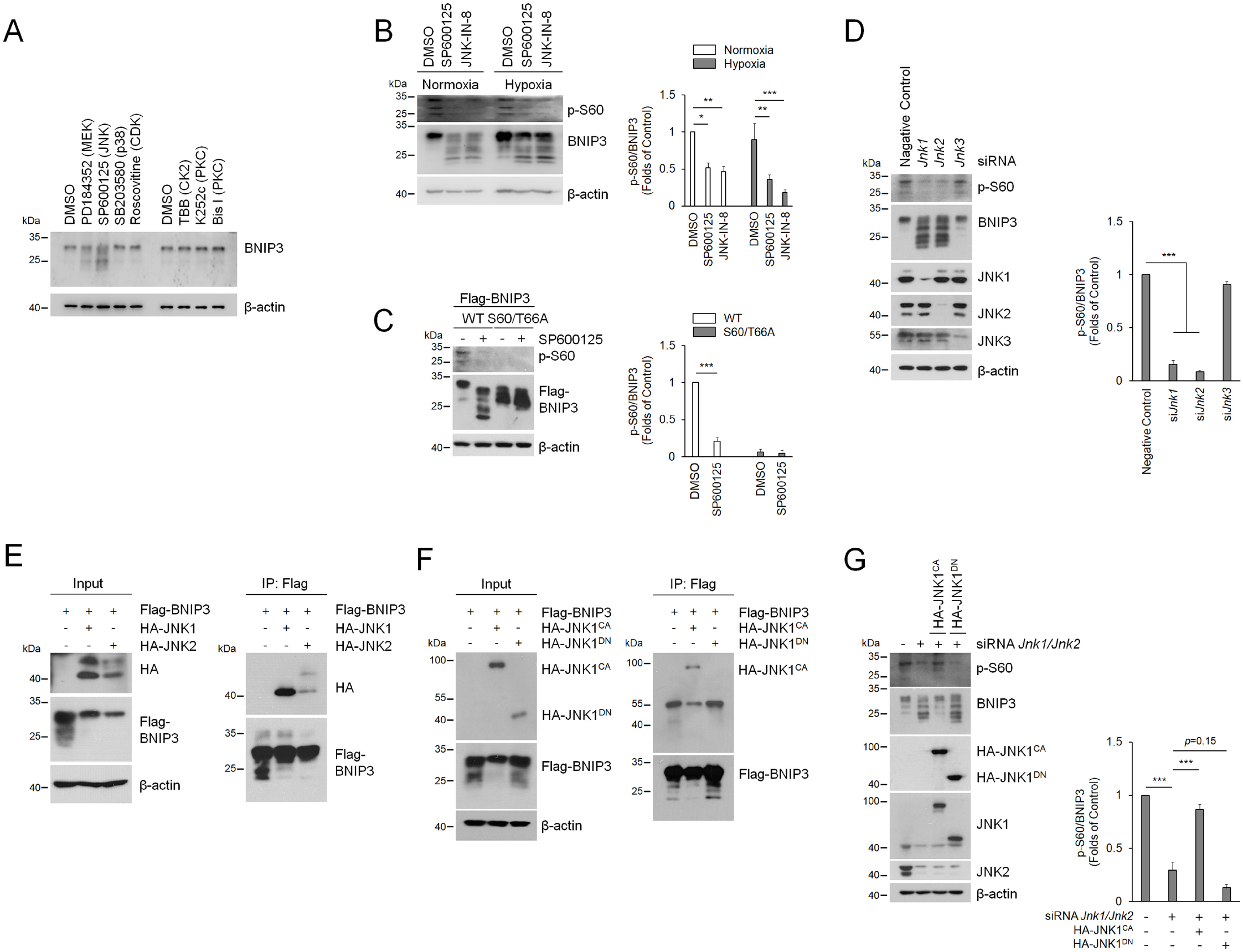
JNK1/2 is the kinase responsible for BNIP3 phosphorylation. A PC12 cells were treated with various kinase inhibitors, including 10 μM PD184352 (MEK inhibitor), 10 μM SP600125 (JNK inhibitor), 10 μM SB203580 (p38 inhibitor), 10 μM Roscovitine (CDK inhibitor), 100 μM TBB (CK2 inhibitor), 10 μM K252c and 1 μM Bis I (PKC inhibitor). Then, cell lysates were subjected to western blotting with the indicated antibodies. DMSO, dimethylsulfoxide. B The phosphorylation of BNIP3 was detected via western blotting after PC12 cells were treated with JNK inhibitor SP600125 (10 μM) or JNK-IN-8 (10 μM) and exposed to 20% O_2_ (Normoxia) or 0.3% O_2_ (Hypoxia) for 6 h. *n* = 3. C PC12 cells stably expressing WT or S60/T66A mutant Flag-BNIP3 were treated with DMSO or SP6000125 and followed by detection of phosphorylation of BNIP3. *n* = 3. D After *Jnk* was knocked down with the indicated siRNA in PC12 cells, the levels of JNK protein and BNIP3 phosphorylation were measured by western blotting. *n* = 3. E, F HeLa cells were transfected with Flag-BNIP3 and HA-JNK1 or HA-JNK2 (E) or constitutively active JNK1 (HA-JNK1CA) (HA-MKK7-JNK1) or dominant negative JNK1 (HA-JNK1^DN^) (HA-JNK1-APF) (F) for 48 h. Cell lysates were then immunoprecipitated with an anti-Flag antibody and detected by western blotting with an anti-HA or anti-Flag antibody. G *Jnk1* and *Jnk2* knockdown PC12 cells were transfected with HA-JNK1CA or HA-JNK1^DN^ mutants for 48 h, cell lysates were then analyzed via western blotting with the indicated antibodies. *n* = 3. The data are expressed as means ± SEM. \**P* < 0.05, \*\**P* < 0.01, \*\*\**P* < 0.001 versus the indicated group.

Given that SP600125 is a broad-spectrum JNK inhibitor for JNK1, JNK2, and JNK3 and PD184352 inhibits ERK1/2 and ERK5 activities, we knocked them down with their respective siRNA to identify the specific BNIP3 kinases. When *Jnk1, Jnk2, Jnk3, Erk1, Erk2* or *Erk5* was separately knocked down with the corresponding siRNA, only *Jnk1* and *Jnk2* knockdown resulted in a significant downshift of BNIP3 bands (Fig. EV3B), which illustrates that JNK1 and JNK2 are the potential kinases for BNIP3 phosphorylation. Subsequently, we confirmed that JNK1 and JNK2, but not JNK3, are the kinases of BNIP3, since knockdown of *Jnk1, Jnk2* but not *Jnk3* directly resulted in a decrease in BNIP3 phosphorylation levels (Fig. 4D).To investigate whether JNK1 and JNK2 interact with BNIP3 in cells, we carried out a co-IP assay after cells were co-transfected with Flag-BNIP3 and HA-JNK1 or HA-JNK2. The results showed that JNK1 had a stronger binding affinity with BNIP3 than JNK2 (Fig. 4E). Hence, we used constitutively active (CA) or dominant negative (DN) JNK1 to further test the interaction of JNK1 with BNIP3. As expected, BNIP3 interacted with CA-JNK1 but not DN-JNK1 (Fig. 4F), demonstrating that the kinase activity of JNK1 is required for the interaction with BNIP3. To elucidate whether Ser 60/Thr 66 of BNIP3 is the phosphorylation site for JNK1, we knocked down *JNK1* and *JNK2* and then overexpressed CA-JNK1 or DN-JNK1 to examine their effects on the phosphorylation of BNIP3 WT or the S60/T66A mutant. The results showed that CA-JNK1 phosphorylated BNIP3 WT instead of the S60/T66A mutant and that DN-JNK1 had no significant effects on phosphorylation (Fig. EV3C), suggesting that Ser 60/Thr 66 of BNIP3 is the phosphorylation site for JNK1. Consistent with this, the effect of JNK1 activity on phosphorylation of BNIP3 was verified by using a phospho-specific antibody (Fig. 4G). Taken together, these data demonstrate that JNK1/2 is a kinase that phosphorylates BNIP3.

### Phosphorylation of BNIP3 S60/T66 by JNK1/2 enhances mitophagy via impeding BNIP3 proteasomal degradation

To investigate the role of JNK1/2-mediated BNIP3 phosphorylation, we first examined the effects of *Jnk1* and *Jnk2* knockdown on BNIP3 phosphorylation and mitophagy. Western blot analysis revealed that knockdown of *Jnk1* and *Jnk2* under both normoxic and hypoxic conditions obviously reduced BNIP3 phosphorylation and simultaneously inhibited mitophagy (Fig. 5A). To further clarify whether phosphorylation of BNIP3 at Ser 60/Thr 66 by JNK1/2 is involved in the regulation of mitophagy, we compared the roles of BNIP3 WT and the S60/T66A mutant in the induction of mitophagy when CA-JNK1 or DN-JNK1 was ectopically overexpressed. Fluorescence images showed that overexpression of CA-JNK1 stimulated the formation of GFP-LC3 puncta and reduced the number of mitochondria in BNIP3 WT cells but not in S60/T66A-expressing cells, while overexpression of DN-JNK1 was ineffective for the induction of mitophagy (Fig. 5B and Fig. EV4). Furthermore, a rescue experiment demonstrated that CA-JNK1 instead of DN-JNK1 restored the BNIP3-LC3 interaction attenuated by *JNK1* and *JNK2* knockdown in WT but not in the S60/T66A mutant cells (Fig. 5C). Taken together, these results consistently demonstrate that JNK1/2 promotes mitophagy by phosphorylating BNIP3 at Ser 60/Thr 66.

**Figure 5.**
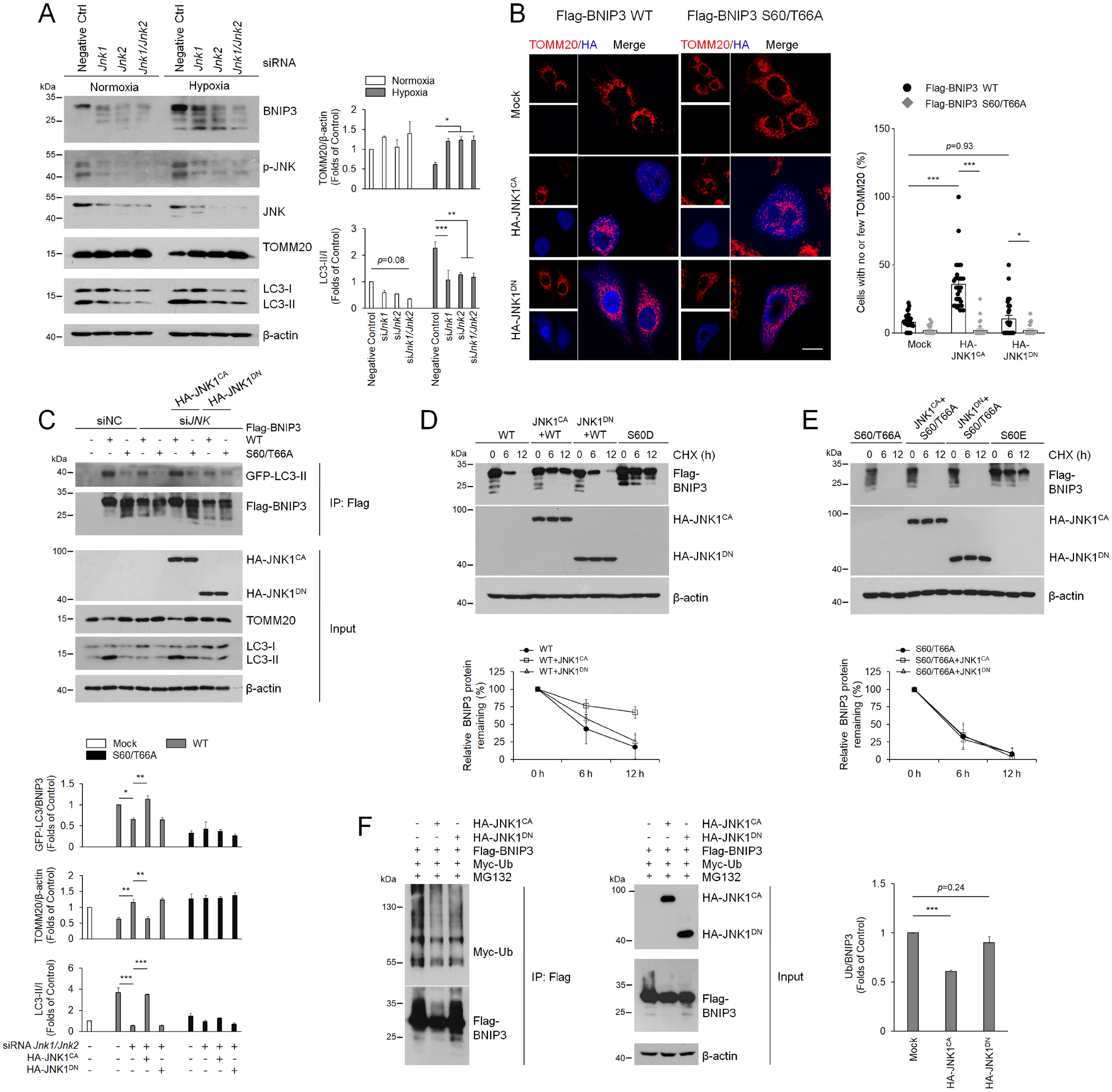
Phosphorylation of BNIP3 at S60/T66 by JNK enhances mitophagy and impedes BNIP3 proteasomal degradation. A PC12 cells were transfected with negative control (NC), *Jnk1* and *Jnk2* siRNA for 48 h, and cell lysates were subjected to western blot analysis with the indicated antibodies. *n* = 3. B Flag-BNIP3 stably expressed HeLa cells that were transfected with plasmids encoding constitutively active (CA) or dominant negative (DN) HA-JNK1 and immunostained with HA (blue) and TOMM20 (Red). The percentage of cells with no or few TOMM20 was quantified. Scale bars, 10 μm. *n* = 30. C HeLa cells were transfected with NC or *JNK1* and *JNK2* siRNA and plasmids encoding GFP-LC3. After 48 h, the cells were transfected with plasmids encoding wild-type (WT) or S60/66A Flag-BNIP3 and HA-JNK1CA or HA-JNK1^DN^ for an additional 24 h. Cell lysates were immunoprecipitated with an anti-Flag antibody and examined via western blotting with the indicated antibodies. *n* = 3. D, E HeLa cells were co-transfected with constitutively active or dominant negative HA-JNK1 and BNIP3 WT (D, left) or the S60/T66A mutants (E, right). After transfection for 48 h, 20 μg ml^-1^ CHX was added to the cultures for the indicated time, and the degradation of BNIP3 was detected via western blotting with the indicated antibodies and quantified, respectively. *n* = 3. F HeLa cells were co-transfected with Flag-BNIP3, Myc-Ub and constitutively active or dominant negative HA-JNK1 for 48 h, and 10 μM MG132 was added 12 h before samples were collected. Cell lysates were boiled and immunoprecipitated with an anti-Flag antibody. The immune complexes were then analyzed via western blotting. *n* = 3. The data are expressed as means ± SEM. \**P* < 0.05, \*\**P* < 0.01, \*\*\**P* < 0.001 versus the indicated group.

In addition to enhancement of the interaction between BNIP3 and LC3 by JNK1/2, we also wondered whether JNK1/2 is directly involved in regulation of BNIP3 stability via phosphorylation of the Ser 60/Thr 66 residue, since the phosphorylation of BNIP3 Ser 60/Thr 66 was shown to improve its stability. Therefore, to determine whether JNK1/2 regulates BNIP3 stability via the Ser 60/Thr 66 site, we examined the effects of JNK1 activity on the stability of BNIP3 after cells were co-transfected with the CA-JNK1 or DN-JNK1 and BNIP3 WT or mutant expression plasmids and then CHX was added to inhibit new protein synthesis. The results clearly showed that compared with BNIP3 WT without JNK1 stimulation, CA-JNK1 increased the stability of BNIP3, similar to that of S60D or S60E, while DN-JNK1 did not affect the stability of BNIP3 WT. On the other hand, neither CA-JNK1 nor DN-JNK1 altered the effect of the phosphorylation-disabled S60/T66A mutant on BNIP3 stability (Fig. 5D, E), indicating that JNK1/2 regulation of BNIP3 stability is achieved by phosphorylation of the Ser 60/Thr 66 residue. To further determine whether JNK1/2 regulation of BNIP3 stability involves the ubiquitin-proteasome pathway, we co-transfected cells with CA-JNK1 or DN-JNK1, BNIP3 and Ub and then conducted a Co-IP assay. We were pleasantly surprised to find that CA-JNK1 significantly reduced the conjugation of ubiquitin to BNIP3 (Fig. 5F). Altogether, our results suggest that JNK1/2 improves the stability of BNIP3 by preventing its degradation via the ubiquitin-proteasome pathway. Thus, we propose that phosphorylation at the Ser 60/Thr 66 residue by JNK1/2 impedes the degradation of BNIP3 via the ubiquitin-proteasome pathway and that the stabilized BNIP3 promotes mitophagy via enhanced interaction with LC3.

### PP1/2A dephosphorylate BNIP3 and suppresses mitophagy by facilitating BNIP3 proteasomal degradation

Since dephosphorylation of BNIP3 is negatively correlated with mitophagy (Fig. 1G and Fig. 2F), we next wondered which protein phosphatase is responsible for dephosphorylation of BNIP3. Given that OA reversed the downward shift of the BNIP3 protein bands (Fig. 1F), which indicates that OA blocks dephosphorylation of BNIP3, and more importantly, OA is a potent inhibitor of the protein phosphatases PP1 and PP2A (Shi, 2009), We therefore speculate that PP1 and PP2A may be the phosphatases of BNIP3. Comparing calyculin A (Cal A) and OA, which are more potent inhibitors of PP1 and PP2A, respectively, we observed both Cal A and OA are all effective in preventing BNIP3 dephosphorylation (Fig. 6A). To determine which catalytic subunit of PP1 or PP2A to play the key role in regulation of BNIP3 dephosphorylation, we found when all catalytic subunits of PP1 or PP2A were knocked down at the same time, the effect of PP1 or PP2A knockdown was manifested, in other words, inhibition of PP1 or PP2A expression increased the level of BNIP3 phosphorylation after cells were exposed to hypoxia instead of normoxia (Fig. 6B). Additionally, we also noticed that the role of PP1 is slightly stronger than PP2A. Therefore, we focused on determining whether PP1 interacts with BNIP3 and affects BNIP3-mediated mitophagy. We co-transfected HeLa cells with PPP1CA or PPP1CC (two main catalytic subunits of PP1) combined with BNIP3 plasmids and observed the effect of PPP1CA/C on BNIP3 phosphorylation and the interaction between them using a co-IP assay. As expected, PPP1CA/C caused remarkable dephosphorylation of BNIP3 when PPP1CA/C was overexpressed in cells. In the meantime, when BNIP3 was immunoprecipitated with a Flag antibody, PPP1CA/C was also pulled down, demonstrating an interaction between PP1 and BNIP3 in cells (Fig. 6C). To examine the effect of PP1 on BNIP3-mediated mitophagy, we transfected cells with BNIP3 and a concentration gradient of PPP1CA/C. Then, we found that with an increase in PPP1CA/C concentration, the phosphorylation of BNIP3 was significantly reduced, and mitophagy was inhibited synchronously (Fig. 6D). These data indicate that PPP1CA/C is a phosphatase that dephosphorylates BNIP3, which largely results in suppression of mitophagy.

**Figure 6.**
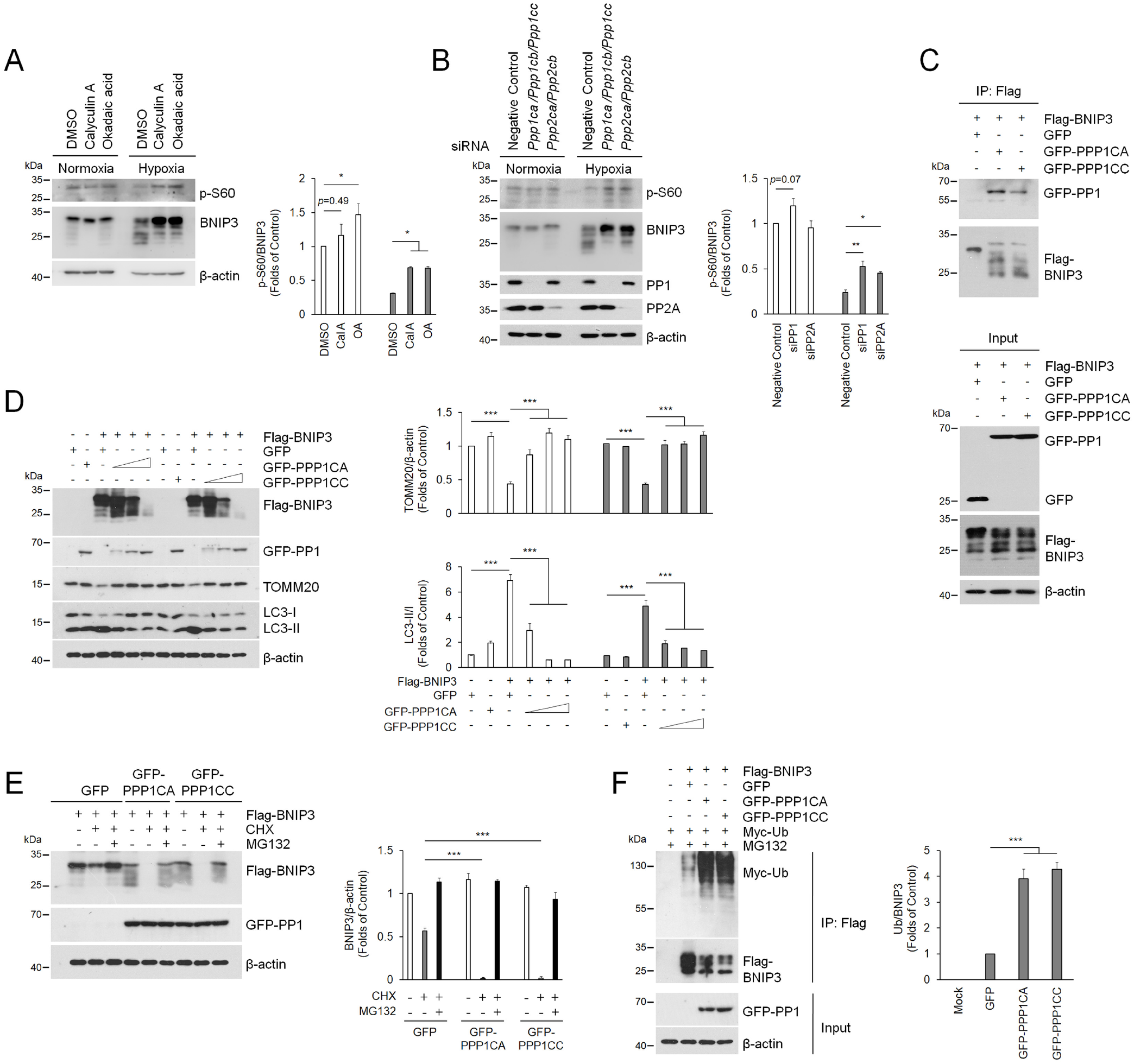
PP1 and PP2A are phosphatases for BNIP3 and suppresses mitophagy by accelerating BNIP3 proteasomal degradation. A PC12 cells were treated with PP1 and PP2A inhibitors, Calyculin A (Cal A, 5 nM) and okadaic acid (OA, 200 nM) and then exposed to 20% O_2_ (Normoxia) or 0.3% O_2_ (Hypoxia) for 24 h, the phosphorylation of BNIP3 was detected via western blotting. *n* = 3. B PP1 and PP2A were knocked down with siRNA targeting their respective catalytic subunits, *Ppp1ca, Ppp1cb, Ppp1cc*, and *Ppp2ca, Ppp2cb*, and then exposed to 20% O_2_ or 0.3% O_2_ for 24 h, after that, western blotting was used to detect the phosphorylation of BNIP3. *n* = 3. C HeLa cells were co-transfected with GFP-Vector, GFP-PPP1CA or GFP-PPP1CC and Flag-BNIP3 for 48 h, and cell lysates were then immunoprecipitated with an anti-Flag antibody and subjected to western blotting with an anti-GFP or anti-Flag antibody. D HeLa cells were transfected with Flag-BNIP3 and GFP-Vector or increasing concentrations of GFP-PPP1CA/PPP1CC plasmids (0.25, 0.5, 1.25 μg ml^-1^) for 48 h, and cell lysates were analyzed by western blotting with the indicated antibodies. *n* = 3. E HeLa cells were co-transfected with Flag-BNIP3 and GFP-Vector, GFP-PPP1CA or GFP-PPP1CC. At 48 h after transfection, the cells were treated with or without 20 μg ml^-1^ CHX or 10 μM MG132 for 12 h. The degradation of BNIP3 was assessed by western blotting with the indicated antibodies. *n* = 3. F HeLa cells were co-transfected with Flag-BNIP3, Myc-Ub and GFP-Vector or GFP-PPP1CA/GFP-PPP1CC for 48 h, and 10 μM MG132 was added 12 h prior to sample collection. Cell lysates were boiled and immunoprecipitated with an anti-Flag antibody. The immune complexes were then analyzed via western blotting. *n* = 3. The data are expressed as means ± SEM. \**P* < 0.05, \*\**P* < 0.01, \*\*\**P* < 0.001 versus the indicated group.

As to how PP1/2A obstructs mitophagy, considering that phosphorylation of BNIP3 improved its stability and promoted mitophagy, we speculated that dephosphorylation of BNIP3 by PP1/2A might negatively regulate the stability of BNIP3, thus leading to failure of mitophagy induction. In addition, we have noticed that the highest concentration of PPP1CA/C leads to nearly complete disappearance of BNIP3, as shown in Fig. 6D, which greatly suggests a negative regulatory effect of PP1 on the stability of BNIP3. Accordingly, we tested the effect of PP1 on the stability of BNIP3 after cells were co-transfected with PPP1CA/C and BNIP3 and then treated with CHX or MG132. We observed an increase in BNIP3 degradation induced by CHX treatment in the pEGFP-C1 control, and the degradation of BNIP3 was exacerbated by PPP1CA/C but blocked by MG132 (Fig. 6E), suggesting that PP1 accelerates BNIP3 degradation via the proteasome pathway. We further demonstrated that PP1 facilitated conjugation of BNIP3 with ubiquitin when cells were co-transfected with PPP1CA or PPP1CC and BNIP3 and Ub plasmids (Fig. 6F), suggesting that PP1 potentiates the degradation of BNIP3 via the ubiquitin-proteasome pathway. Collectively, our results demonstrate that PP1/2A is a phosphatase of BNIP3 and suppresses BNIP3-mediated mitophagy, primarily due to accelerated BNIP3 degradation via the ubiquitin-proteasome pathway.

## Discussion

Regulation of mitophagy under hypoxia is critical for cell fate and is related to whether cells adapt to hypoxia. BNIP3 is a mitophagy receptor that mediates mitophagy and is also a hypoxia-responsive protein that is highly upregulated under hypoxia, but how BNIP3 regulates mitophagy under hypoxia remains a challenging question. The key issue is that the fine regulation of BNIP3 under hypoxia, which involves the upstream modulation of BNIP3-mediated mitophagy, has not been uncovered. It has been reported that overexpression of BNIP3 in hypoxia promotes its interaction with BCL-2 or BCL-XL, which leads to the release of Beclin-1 from BCL-2/BCL-XL binding and hence to autophagy initiation (Bellot *et al*., 2009; Zhang *et al*., 2008). Some studies have also noted that phosphorylation of BCL-2/BCL-XL by JNK promotes the disassociation of Beclin-1 from BCL-2/BCL-XL (Klein *et al*, 2015; Wei *et al*, 2008a; Wei *et al*, 2008b; Zhong *et al*, 2017). However, following autophagy initiation in the same context, how damaged or unwanted mitochondria are recognized, thereby inducing mitophagy, has not been elucidated. Moreover, although phosphorylation of BNIP3 at Ser 17/24 has been shown to enhance its binding to LC3 and promote mitophagy (Zhu *et al.*, 2013), thus far, no kinase or phosphatase that targets BNIP3 Ser 17/24 has been found.

In this study, we first found that overexpression of BNIP3 under hypoxia is not the unique key to determining mitophagy. Because even when BNIP3 was overexpressed in cells exposed to 0.3% O_2_ for more than 12 h, mitophagy was still suppressed (Fig. 1G). Instead, we demonstrated that phosphorylation of BNIP3 is required for the induction of mitophagy, which facilitates cell survival under hypoxia. Subsequently, we identified Ser 60/Thr 66 as a new phosphorylation site in BNIP3. To our surprise, phosphorylation of BNIP3 at Ser 60/Thr 66 is not only essential for mitophagy induction but also for the stability of BNIP3. Our results further show that BNIP3 phosphorylation at the Ser 60/Thr 66 residue hinders its degradation via the ubiquitin-proteasome pathway. Based on these data, we believe that inhibition of BNIP3 proteasomal degradation favors its mediation of mitophagy. In other words, induction of mitophagy is hampered by BNIP3 degradation, similar to the result of *Bnip3* knockdown with siRNA. Overall, mitophagy is largely dependent on the phosphorylation of BNIP3 at Ser 60/Thr 66.

We next identified JNK1/2 as the kinase of BNIP3 at Ser 60/Thr 66, which contributes to induction of mitophagy by enhancing the BNIP3-LC3 interaction. Since JNK also activates BCL-2, which facilitates the dissociation of Beclin-1 from BCL-2 (Wei *et al.*, 2008a; Wei *et al.*, 2008b), we explored whether phosphorylation at Ser 60/Thr 66 by JNK1/2 is involved in the interaction of BNIP3 with BCL-2. Unfortunately, we found that phosphorylation of BNIP3 at Ser 60/Thr 66 is not associated with binding of BCL-2, as the S60/T66A mutant did not alter the interaction of BNIP3 with BCL-2 (Fig. EV2B). However, this result does not rule out the possibility that other phosphorylation sites participate in the binding of BNIP3 to BCL-2. Additionally, we identified PP1and PP2A as BNIP3 phosphatases, which block mitophagy by dephosphorylation of BNIP3. Dephosphorylation of BNIP3 by PP1/2A not only plays a negative regulatory role in the induction of mitophagy but also in the stability of BNIP3. Especially, the degradation of BNIP3 through the ubiquitin-proteasome pathway was significantly accelerated by dephosphorylation. It is a pity that we did not find an E3 ligase for BNIP3 that regulates its proteasomal degradation. However, we can still speculate on such a possibility that phosphorylation at the Ser 60/Thr 66 may block E3-mediated conjugation of ubiquitin to BNIP3, while dephosphorylation at the Ser 60/Thr 66 contributes to the ubiquitin conjugation and subsequent proteasomal degradation of BNIP3. In any case, we propose the hypothesis that BNIP3 phosphorylation at Ser 60/Thr 66 by activated JNK1/2 under moderate hypoxia conditions (10% O_2_ or early stages of 0.3% O_2_) blocks BNIP3 degradation via the ubiquitin-proteasome pathway, and then the stabilized and activated BNIP3 recruits LC3, thereby promoting mitophagy and cell survival; however, PP1/2A activated under severe hypoxia conditions (late stages of 0.3% O_2_) accelerates BNIP3 proteasomal degradation, which impairs the induction of mitophagy and causes cell death (Fig. 7).

**Figure 7.**
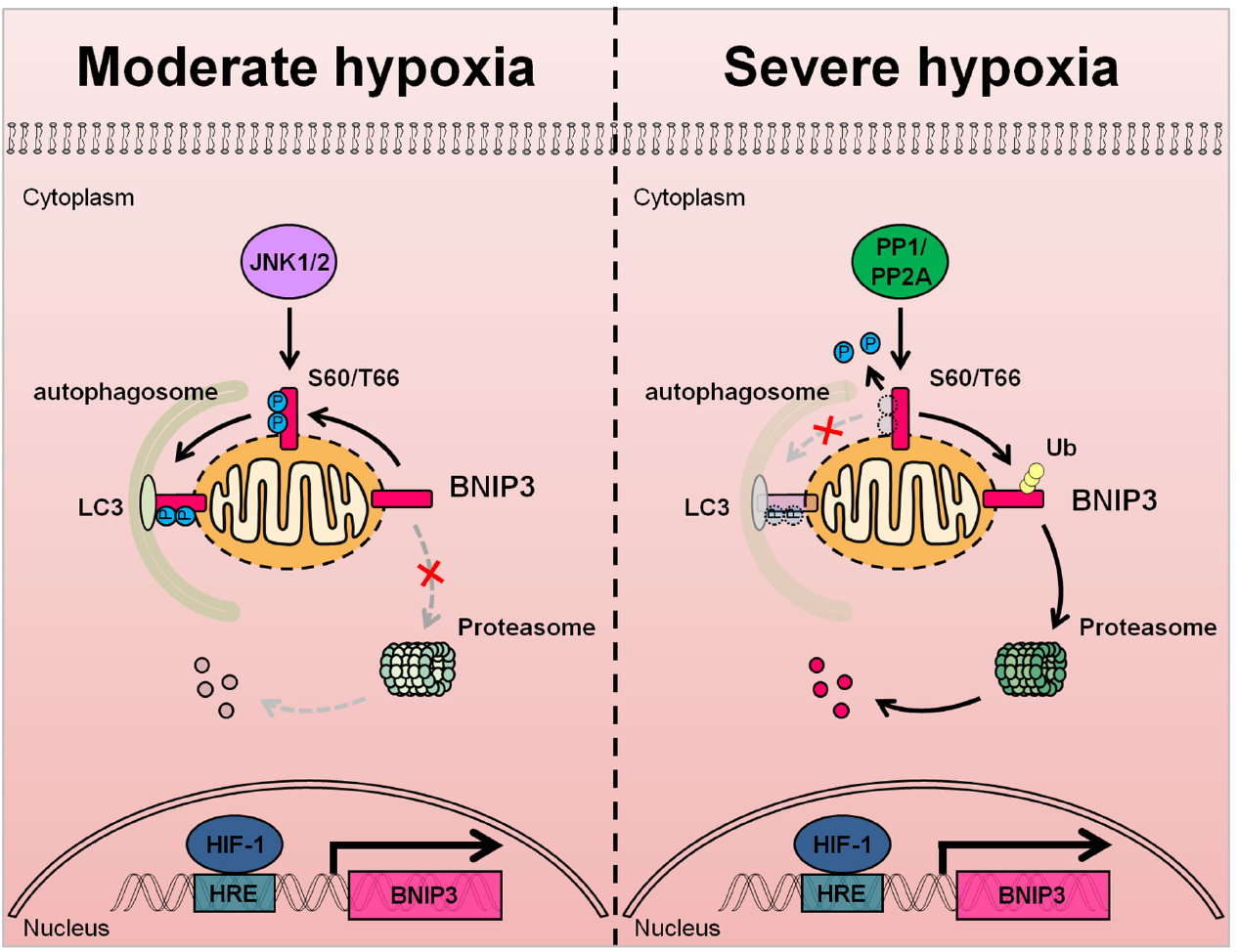
The hypothetical mechanism of BNIP3 phosphorylation-mediated mitophagy under hypoxia. In response to moderate hypoxia (10% O_2_ or early stages of 0.3% O_2_), a generous amount of BNIP3 is phosphorylated at S60/T66 by JNK1/2, which blocks the conjugation of ubiquitin (Ub) to BNIP3, inducing mitophagy activation. In severe hypoxia (late stages of 0.3% O_2_), JNK1/2 is inactivated and BNIP3 is dephosphorylated by PP1 or PP2A, which leads to the recruitment of ubiquitin to BNIP3 and its degradation via the ubiquitin-proteasome pathway, suppressing the induction of mitophagy.

In conclusion, we provide evidence that BNIP3 phosphorylation level is more critical for the induction of mitophagy than its total protein level, and that JNK1/2/ and PP1/2A oppositely regulate the phosphorylation and stability of BNIP3 in response to different hypoxia. Future identification of an E3 ligase for BNIP3 and its regulation under hypoxia will be of great significance for revealing the comprehensive functions of BNIP3. Further study on the biological significance of coordinated regulation of BNIP3 by JNK1/2 and PP1/2A under hypoxic conditions could provide insight into therapeutic strategies against hypoxia-related diseases.

## Materials and Methods

### Cell culture and hypoxia treatment

PC12, HeLa and HEK293T cells were obtained from American Type Culture Collection (ATCC). Cells were cultured in Dulbecco’s modified Eagle’s medium (DMEM, HyClone, SH30081) supplemented with 1% penicillin-streptomycin (HyClone, SV30010) and 5% fetal bovine serum, 10% horse serum (Gibco) or 10% fetal bovine serum at 37 °C under 5% CO_2_. PC12 and Hela cells stably expressing Flag-BNIP3 were selected in media containing 1 μg ml^-1^ puromycin. For hypoxia treatment, cells were placed in an incubator (Thermo Fisher Scientific) at 37 °C with 20% O_2_ and 5% CO_2_ for 24 h, and then moved to a hypoxia chamber (0.3% O_2_, 5% CO_2_ and 94.7% N_2_ (Coy laboratory) or 10% O_2_, 5% CO_2_ and 85% N_2_ (Thermo Fisher Scientific).

### Reagents and antibodies

Mitotracker (M7512) was purchased from Thermo Fisher Scientific; λ-PPase (P0753) was obtained from New England Biolabs; 3-MA (M9281), cycloheximide (01810), K252c (S3939), MG132 (M7449), okadaic acid (O8010) and TBB (T0826) were from Sigma-Aldrich; Bis I (S7208), JNK-IN-8 (S4901), PD184352 (S1020), Roscovtine (S1153), SB203580 (S1076) and SP600125 (S1460) were from Selleck; Calyculin A (1336) was purchased from Tocris.

The following antibodies were used for western blotting: anti-BNIP3 (1:1,000, mouse mAb, Abcam, ab10433), anti-p62 (1:10,000, mouse mAb, Abcam, ab56416), anti-phospho-BNIP3 (Ser 60, 1:400, rabbit pAb) was generated by Abclonal, anti-BCL-2 (1:1,000, mouse mAb, BD bioscience, 610538), anti-JNK1 (1:1,000, mouse mAb, Cell Signaling Technology, 3708), anti-JNK2 (1:1,000, rabbit mAb, Cell Signaling Technology, 9258), anti-JNK3 (1:500, rabbit mAb, Cell Signaling Technology, 2305), anti-PP2A (1:1,000, rabbit pAb, Cell Signaling Technology, 2038), anti-p-MAPK/CDK substrates (1:500, rabbit mAb, Cell Signaling Technology, 2325), anti-p-SAPK/JNK (1:1,000, rabbit pAb, Cell Signaling Technology, 9251), anti-SAPK/JNK (1:1,000, rabbit pAb, Cell Signaling Technology, 9252), anti-Flag (anti-DDDDK, 1:10,000, mouse mAb, MBL, M185-3), anti-Flag (anti-DDDDK, 1:2,000, rabbit pAb, MBL, PM020), anti-HIF-1α (1:1,000, mouse mAb, Novus Biologicals, NB100-105), anti-c-Myc (1:1,000, rabbit pAb, Santa Cruz, sc-789), anti-GFP (1:1,000, rabbit pAb, Santa Cruz, sc-8334), anti-HA (1:1,000, rabbit pAb, Santa Cruz, sc-805), anti-PP1 (1:1,000, mouse mAb, Santa Cruz, sc-7482), anti-TOMM20 (1:5,000, rabbit pAb, Santa Cruz, sc-11415), anti-LC3B (1:4,000, rabbit pAb, Sigma-Aldrich, L7543) and anti-β-actin (1:10,000, mouse mAb, Sigma-Aldrich, A5316). The following HRP-conjugated secondary antibodies were used for western blotting: goat anti-mouse IgG (1:2,000, MBL, 330) and goat anti-rabbit IgG (1:2,000, MBL, 458). The following antibodies were used for immunofluorescence experiments: anti-Flag (anti-DDDDK, 1:1,000, mouse mAb, MBL, M185-3), anti-Flag (anti-DDDDK, 1:1,000, rabbit pAb, MBL, PM020), anti-HA (1:1,000, mouse mAb, MBL, M180-3) and anti-TOMM20 (1:1,000, rabbit pAb, Santa Cruz, sc-11415). The fluorescent secondary antibodies were conjugated with either Alexa Fluor 594 (1:1,000, anti-rabbit, Cell Signaling Technology, 8889) or Alexa Fluor 647 (1:1,000, anti-mouse, Cell Signaling Technology, 4410). Mouse IgG (C2118) was purchased from Applygene.

### Plasmids, transfection and virus production

BNIP3 was amplified from rat cDNA (NCBI RefSeq NM_053420.3) via PCR and fused with Flag via an N-terminal epitope tag. Then, Flag-tagged BNIP3 was cloned into a pcDNA3.1 vector (Thermo Fisher Scientific) or a pCDH vector (System Biosciences, CD550A-1). JNK1 (NCBI RefSeq NM_001323302.1) and JNK2 (NCBI RefSeq NM_002752.4) were amplified from human cDNA via PCR and cloned into a pXJ40-HA vector. Site-directed mutants and siRNA-resistant constructs were performed using kits according to standard methods (SBS Genetech). The primer information can be found in Table S1. All the plasmids were verified by DNA sequencing. pXJ40-HA, pXJ40-Myc-Ub, pEGFP-C1-PPP1CA, pEGFP-C1-PPP1CC and pEGFP-C1-LC3B (Xu *et al*, 2018) were gifts from Q. Xia and T. Zhou (State Key Laboratory of Proteomics, Beijing). pSRα-HA-MKK7-JNK1 (JNK1CA) and pSRα-HA-JNK1-APF (JNK1^DN^) (Thr-Pro-Tyr replaced with Ala-Pro-Phe) (Wang *et al*, 2011) were gifts from J-Y. Zhang (Institute of Cognition and Brain Sciences, Beijing).

Transfection of plasmids was performed using Lipofectamine 2000 reagent (Thermo Fisher Scientific) according to the manufacturer’s instructions. For RNA interference, cells were transfected with negative control or with predesigned siRNAs (Sigma-Aldrich or Ribobio) targeting the indicated genes (Table S2) at a final concentration of 50 nM or 100 nM using X-tremeGene siRNA transfection reagent (Roche, 04476093001) according to the manufacturer’s instructions. For lentivirus production, pCDH-Flag-BNIP3 was co-transfected with psPAX2 and pMD2.G plasmids into HEK293T cells.

### Transmission electron microscopy

After PC12 cells were exposed to different oxygen concentrations for 24 h, the cells were collected by digestion and centrifugation, washed with PBS, and then fixed with 3% glutaraldehyde in 0.075 M PBS (pH 7.4) at 4 °C for 2 h. After fixation, the cells were washed three times with PBS, post-fixed in 1% OsO_4_ at 4 °C for 1 h, and washed in PBS for 15 min. Cell precipitates were dehydrated through a graded series of 50% to 90% ethanol, which was then replaced with 90% to 100% acetone. The cultures were soaked and then embedded in acrylic resin. Next, 60-nm ultrathin sections were collected on copper grids and stained with uranyl acetate for 10 min and lead citrate for 10 min. The samples were visualized and photographed using a HITACHI H-7650 transmission electron microscope at 80 kV.

### Confocal imaging of living cells

For mitophagy analysis, PC12 cells were transfected with plasmid encoding GFP-LC3 for 48 h and then exposed to hypoxia or normoxia for an additional 24 h. Next, mitochondria were marked with 50 nM MitoTracker Red CMXRos at 37°C under 5% CO_2_ for 30 min, and then, the cells were washed with PBS. Cell images were captured with a confocal microscope (Carl Zeiss). Autophagosome formation and mitochondrial co-localization were analyzed using ImageJ software (NIH). The co-localization of autophagosome and mitochondria was quantified by counting more than 40 cells.

### Immunofluorescence microscopy

HeLa cells were grown on glass coverslips. After the indicated treatment, cells were fixed with 4% paraformaldehyde (PFA) for 15 min at room temperature, washed three times with PBS, permeabilized with 0.5% Triton X-100 in PBS for 30 min and blocked with 5% goat serum for 1 h at room temperature. Cells were then incubated with primary antibodies diluted in goat serum overnight at 4°C, followed by incubation with secondary antibodies diluted in goat serum for 1 h. All images were captured with a confocal microscope. For quantification of no or few TOMM20, more than 150 cells from 30 different fields were counted.

### Real-time quantitative PCR

Total RNA was isolated from cells using Trizol^®^ reagent (Thermo Fisher Scientific, 15596-026) according to the manufacturer’s protocol. An aliquot of 1 μg of total mRNA was reversely transcribed at 42°C for 1 h in a 10 μl reaction mixture containing oligo (dT) 18 primer, M-MLV reverse transcriptase and RNase inhibitor (TaKaRa, D2639A). Q-PCR was performed in triplicate with Power SYBR^®^ Green (Thermo Fisher Scientific, 4367659) on a Step-one Plus system (Thermo Fisher Scientific). The primer information can be found in Table S3.

### SDS-PAGE and western blotting

Cells were lysed in ice-cold RIPA buffer (50 mM Tris-HCl (pH 7.4), 150 mM NaCl, 1% NP-40, and 0.1% SDS) containing a protease inhibitor cocktail (Roche, 11697498001). Equal amounts of protein were separated on 10%-15% SDS-PAGE gels and transferred to PVDF membranes (Roche, 03010040001). The membranes were probed with the indicated primary antibodies followed by the appropriate HRP-conjugated secondary antibodies. The protein content was determined with a chemiluminescence (ECL) assay kit (Bio-Rad, 1705060).

### Lambda phosphatase assay

PC12 cells were seeded in 60-mm dishes. After the indicated treatment, cells were washed with ice-cold PBS and lysed in RIPA buffer containing protease inhibitor cocktail. After centrifugation at 13,200 g for 15 min at 4°C, supernatant fractions containing equal amounts of protein (50 μg) were incubated with lambda phosphatase and the phosphatase buffer for 1 h at 37°C. Samples were boiled and subjected to SDS-PAGE and western blotting.

### Immunoprecipitation

After transfection with the indicated plasmids, HeLa cells were collected and lysed in 1 ml of lysis buffer (20 mM Tris-HCl, 150 mM NaCl, 1 mM Na_2_-EDTA, 1 mM EGTA, 1% Triton X-100, 2.5 mM sodium pyrophosphate, 1 mM β-glycerophosphate, 1 mM Na_3_VO_4_, and 1 μg ml^-1^ leupeptin, Cell Signaling Technology, 9803) containing protease inhibitor cocktail. After centrifugation at 13,200 g for 15 min at 4°C, equal amounts of cell lysates were preincubated with mouse IgG (Applygene, C2118) and Protein A/G agarose beads (Santa Cruz, sc-2003) for 4 h at 4°C and then clarified by centrifugation. Supernatant fractions were immunoprecipitated with 2 μg anti-Flag antibody (MBL, M185-3) and 40 μl Protein A/G agarose beads overnight at 4°C. After being washed five times, the immunoprecipitates were boiled in Laemmli sample buffer (Bio-Rad, 1610737) for 5 min. Samples were analyzed via SDS-PAGE and western blotting.

### Degradation and ubiquitylation assays

BNIP3 degradation was estimated using cycloheximide (CHX) chase assays. Cells were treated with 20 μg ml^-1^ CHX for the indicated time, and cell lysates were subjected to SDS-PAGE and western blotting. For the ubiquitination assay, cells were lysed in RIPA buffer containing protease inhibitor cocktail and boiled for 10 min. After being clarified by centrifugation, cell lysates were precleared with mouse IgG and immunoprecipitated with an anti-Flag antibody and Protein A/G agarose beads. Immunoprecipitated Flag-BNIP3 was detected via SDS-PAGE and western blotting.

### Statistical analysis

The statistical data are expressed as the mean ± SEM. Statistical significant differences were assessed using oneway analysis of variance (ANOVA) followed by Tukey’s multiple comparison test (Fig. 1B, Fig. 1C, Fig. 2D, Fig. 2E, Fig. 2G, Fig. 2H, Fig. 3G, Fig. 4D, Fig. 5F, Fig. 6F, Fig. EV1A, Fig. EV2A, Fig. EV2B) or Dunnett’s multiple comparison test (Fig. 1G, Fig. EV1B) or two-way ANOVA with Tukey’s multiple comparison test (Fig. 1D, Fig. 2F, Fig. 4B, Fig. 4C, Fig. 4G, Fig. 5A, Fig. 5B, Fig. 5C, Fig. 6A, Fig. 6B, Fig. 6D, Fig. 6E, Fig. EV3C). Differences between compared groups were considered statistically significant at *P* values < 0.05. \**P*<0.05, \*\**P* < 0.01 and \*\*\**P* < 0.001 versus the corresponding controls are indicated. All statistical analyses were done using GraphPad Prism 8 software.

## Acknowledgement

We thank Prof. Q. Xia and T. Zhou (State Key Laboratory of Proteomics, Beijing) for providing the pXJ40-HA, pXJ40-Myc-Ub, pEGFP-C1-LC3B, pEGFP-C1-PPP1CA, and pEGFP-C1-PPP1CC plasmids and Prof. J-Y. Zhang (Institute of Cognition and Brain Sciences, Beijing) for providing the pSRα-HA-MKK7-JNK1 (JNK1CA) and pSRα-HA-JNK1-APF (JNK1^DN^) plasmids.This work was supported by grants from the National Natural Science Foundation of China (31271211, 31771321 and 81971781 to L.-Y. Wu), the Key Program of National Nature Sciences Foundation of China (81430044 to L.-L. Zhu) and the National Basic Research Program of China (2011CB910800 to L.-L. Zhu and 2012CB518200 to M. Fan).

## Author contributions

L.-Y. Wu, L.-L. Zhu and M. Fan supervised the project. L.-Y. Wu and Y.-L. He designed and performed the majority of the experiments. S.-H. Gong and M. Zhao contributed to the immunofluorescence analyses. X. Chen and T. Zhao helped to prepare cell lines and reagents. Y.-Q. Zhao provided technical support. L.-Y. Wu and Y.-L. He interpreted the data and wrote the manuscript, with the help of all the authors.

## Supporting Information

### Expanded View Figure legends

**Figure EV1. Hypoxia induces mitophagy and phosphorylation of BNIP3.**

A PC12 cells were exposed to 20% O_2_, 10% O_2_ or 0.3% O_2_ for 24 h. Cells were identified by immunofluorescence staining with antibodies against β-actin (white outline) and TOMM20 (red), and nuclear DNA was marked using DAPI (blue). Scale bars, 10 μm. The percentage of cells with no or few TOMM20 are quantified. *n* = 3.

B Hela cells were exposed to 0.3% O_2_ for the indicated time, the levels of BNIP3 and mitophagy related proteins were detected via western blot. The data are expressed as means ± SEM. \**P* < 0.05, \*\**P* < 0.01, \*\*\**P* < 0.001 versus the indicated group.

Figure EV2. **Phosphorylation of BNIP3 at S60/T66 is not necessary for its binding to BCL-2.**

A HeLa cells were transfected with empty vector, WT or the indicated mutated Flag-BNIP3, and then, the cell lysates were subjected to immunoprecipitation (IP) with an anti-Flag antibody. Phosphorylation of BNIP3 at S60/T66 was detected in the immune complexes via western blotting using an anti-phospho-MAPK/CDK substrate antibody. *n* = 3.

B HeLa cells were transfected with empty vector, WT or the indicated Flag-BNIP3 mutants for 48 h. Cell lysates were immunoprecipitated with the anti-Flag antibody and then subjected to western blot analysis with BCL-2 and Flag antibodies. *n* = 3. The data are expressed as means ± SEM. \**P* < 0.05, \*\**P* < 0.01, \*\*\**P* < 0.001 versus the indicated group.

**Figure EV3. JNK1/2 is the kinase responsible for BNIP3 phosphorylation.**

A PC12 cells were treated with okadaic acid (OA) and PD184352 (left) or SP600125 (right) for 12 h and analyzed via western blotting with an anti-BNIP3 or anti-β-actin antibody.

B PC12 cells were transfected with negative control (NC) or the indicated siRNA for 48 h, and the mRNA levels of related genes (left) and BNIP3 expression (right) were detected by real-time PCR and western blotting, respectively. *n* = 9.

C *JNK1* and *JNK2* knockdown Hela cells were transfected with WT or S60/T66A and HA-JNK1CA or HA-JNK1^DN^ mutants, and 48 h post-transfection, cell lysates were immunoprecipitated with an anti-Flag antibody. The immune complexes were then analyzed via western blotting with the indicated antibodies. *n* = 3. The data are expressed as means ± SEM. *P < 0.05, **P < 0.01, ***P < 0.001 versus the indicated group.

**Figure EV4. BNIP3 phosphorylation at S60/T66 by JNK1 enhances mitophagy.** Representative images of GFP-LC3 puncta in Flag-BNIP3 stable stably expressed HeLa cells, co-transfected with plasmids encoding HA-JNK1. Cells were identified by immunofluorescence staining with antibodies against HA (blue) and Flag (red). Scale bars, 10 μm.

## Appendix

**Table S1.**
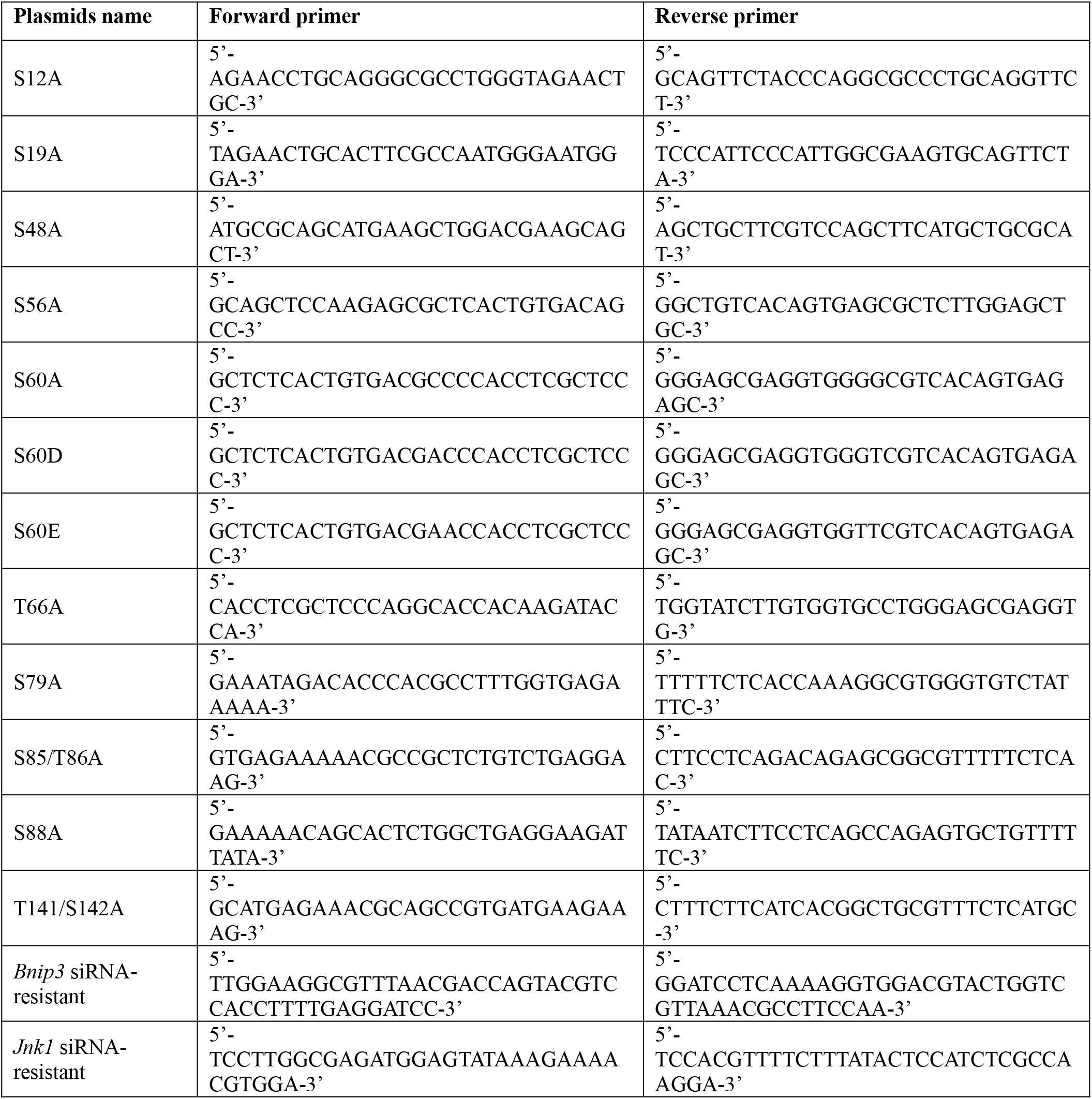
The primer information for the site-directed mutagenesis and the siRNA-resistant constructs.

**Table S2.**
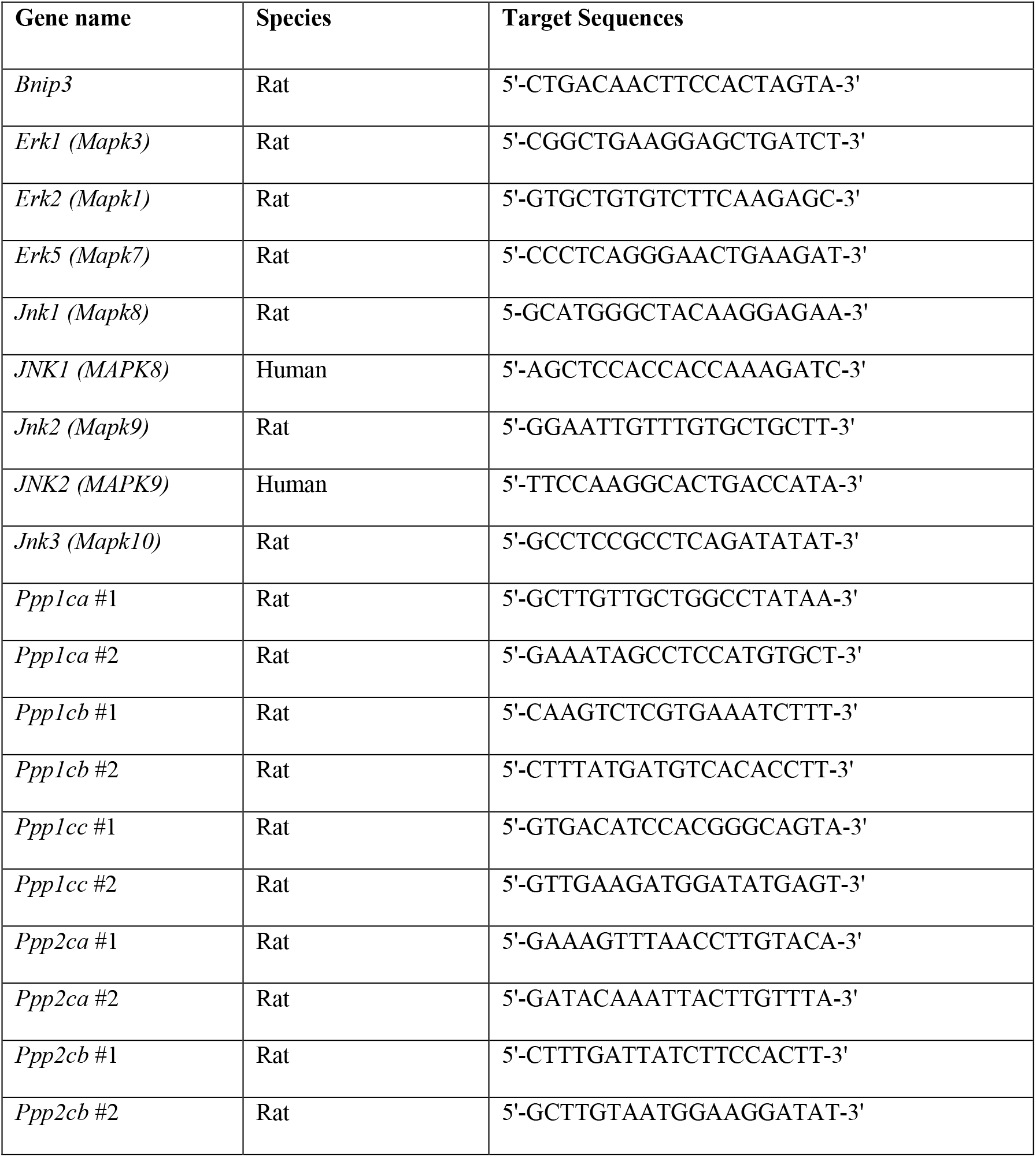
List of siRNA used in this study.

**Table S3.**
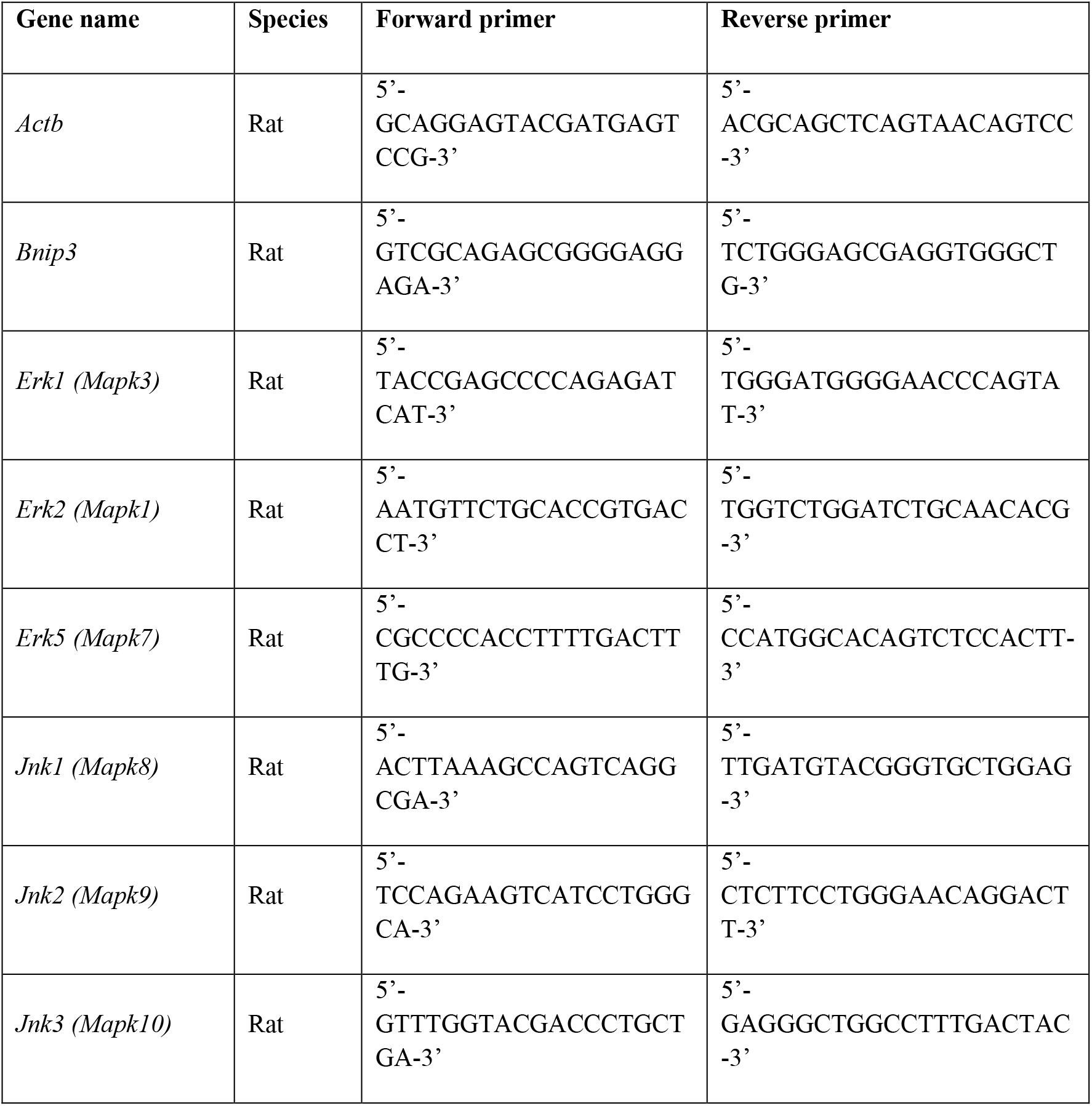
Sequence of real-time PCR primer used in this study.

